# Uncovering cryptic pockets in the SARS-CoV-2 spike glycoprotein

**DOI:** 10.1101/2021.05.05.442536

**Authors:** Lorena Zuzic, Firdaus Samsudin, Aishwary T. Shivgan, Palur V. Raghuvamsi, Jan K Marzinek, Alister Boags, Conrado Pedebos, Nikhil Kumar Tulsian, Jim Warwicker, Paul MacAry, Max Crispin, Syma Khalid, Ganesh S. Anand, Peter J Bond

**Author notes:** Joint first authors (Lorena Zuzic, Firdaus Samsudin).

## Abstract

The recent global COVID-19 pandemic has prompted a rapid response in terms of vaccine and drug development targeting the viral pathogen, severe acute respiratory syndrome coronavirus 2 (SARS-CoV-2). In this work, we modelled a complete membrane-embedded SARS-CoV-2 spike (S) protein, the primary target of vaccine and therapeutics development, based on available structural data and known glycan content. We then used molecular dynamics (MD) simulations to study the system in the presence of benzene probes designed to enhance discovery of cryptic, potentially druggable pockets on the S protein surface. We uncovered a novel cryptic pocket with promising druggable properties located underneath the 617-628 loop, which was shown to be involved in the formation of S protein multimers on the viral surface. A marked multi-conformational behaviour of this loop in simulations was validated using hydrogen-deuterium exchange mass spectrometry (HDX-MS) experiments, supportive of opening and closing dynamics. Interestingly, the pocket is also the site of the D614G mutation, known to be important for SARS-CoV-2 fitness, and within close proximity to mutations in the novel SARS-CoV-2 strains B.1.1.7 and B.1.1.28, both of which are associated with increased transmissibility and severity of infection. The pocket was present in systems emulating both immature and mature glycosylation states, suggesting its druggability may not be dependent upon the stage of virus maturation. Overall, the predominantly hydrophobic nature of the cryptic pocket, its well conserved surface, and proximity to regions of functional relevance in viral assembly and fitness are all promising indicators of its potential for therapeutic targeting. Our method also successfully recapitulated hydrophobic pockets in the receptor binding domain and N-terminal domain associated with detergent or lipid binding in prior cryo-electron microscopy (cryo-EM) studies. Collectively, this work highlights the utility of the benzene mapping approach in uncovering potential druggable sites on the surface of SARS-CoV-2 targets.

---

The rapidly spreading outbreak of COVID-19 caused by a novel coronavirus, SARS-CoV-2^1^, has triggered an unprecedented scale of global socioeconomic meltdown.^2^ Central to the mechanism of infection is the S protein on the surface of the virion, which is the primary target for vaccine and therapeutics development. The S protein is a class 1 viral fusion protein trimer composed of two major subunits: S1, which facilitates host cell recognition by interacting with the human angiotensin converting enzyme 2 (ACE2), and S2, which mediates membrane fusion and entry into the host cell. To date, several structures of the prefusion S protein ectodomain (ECD) and its receptor binding domain (RBD) bound to ACE2 have been resolved using cryo-EM and X-ray crystallography.^3–8^ The S protein ECD is a large trimeric protein made of various functioning domains and predominantly covered with glycans.^9^

Structural and biophysical studies along with MD simulations have shown that the S protein is highly dynamic.^10–13^ Crucially, recent cryo-EM structures have uncovered cryptic pockets in the RBD and N-terminal domain (NTD), which serve as potential druggable epitopes.^14–16^ In recent years, simulations of therapeutically relevant proteins with small organic probes have successfully been used to induce energetically unfavourable opening of hydrophobic cryptic pockets and subsequently identify novel druggable sites.^17–19^ In this work, we thus built a membrane-bound glycosylated model of the S protein and simulated it in the presence of a solution containing benzene probes, to enhance the sampling of novel cryptic pockets that could potentially be targeted by small molecules, peptides, or monoclonal antibodies.

The S protein RBD defines its predominant functional state, with the open state having one RBD in the “up” conformation, allowing for binding to the host ACE2 receptor,^8,20^ with the other two RBDs in the “down” conformation interacting with NTD and other subdomains. We first built a complete model of the SARS-CoV-2 S protein in this open “up-down-down” RBD configuration (see Supplementary Methods). The effects of glycosylation on S protein dynamics were investigated by generating three different glycoforms: i) with the most dominant species of glycans based on mass spectrometry data;^9^ ii) with only high-mannose glycans to represent the unprocessed glycoform; and iii) without glycans. To simulate the S protein in its native membrane environment,^22,23^ we embedded our models in an endoplasmic reticulum-Golgi intermediate compartment (ERGIC) membrane. A series of 200 ns simulations were then carried out in the presence or absence of 0.2 M benzene (see Table S1 for a complete list of simulations).

Interestingly, in all simulations the S protein did not maintain an upright conformation with respect to the plane of the membrane. Instead, we observed a tilting motion of the ECD of up to 90°, facilitated by the two flexible hinges between the ECD and HR2 domains as well as between the HR2 and transmembrane (TM) domains (Figure S1). The flexible bending motion results in orientational freedom of the ECD, presumably yielding a more expansive sampling of the RBD at the host cell surface. This may potentially increase the probability of binding to the ACE2 receptor and hence contribute towards efficient virus-host cell recognition. Such structural dynamics are in good agreement with a range of experimental data including from HDX-MS,^10^ cryo-EM of recombinant S protein ECD,^3^ and cryo-electron tomography (cryo-ET) of intact SARS-CoV-2 virions,^12,13,24^ as well as simulation studies of independently built S protein models.11,25 As expected, in simulations of the S protein modelled with either the predominant glycan species or oligomannose-type glycans, the glycans showed a high degree of mobility resulting in a larger surface area of the S protein being covered by them compared to what is observed in static structures (Figure S2). The glycans covered a larger percentage of the stalk surface compared to the ECD, in agreement with previous simulations.^11,25^ Collectively, the observed protein and glycan dynamics of our S glycoprotein models thus correlate well with other independent experimental and computational studies.

We next set out to uncover cryptic binding pockets on the surface of the S glycoprotein that could potentially represent targets for rational drug design. Hence, we performed a series of 200 ns simulations of the membrane-embedded, full-length S protein models with a 0.2 M concentration of benzene molecules within the bulk solvent. The benzene parameters have been modified to prevent accumulation within the hydrophobic lipid environment (details in Supplementary Methods).^26^ This was confirmed by the low percentage of benzene found in contact with membrane lipids throughout the simulations, and a similar progression of area per lipid and membrane thickness compared to simulations without benzene, indicating that the presence of benzene did not alter the membrane environment (Figure S3). The stable secondary structure of the whole S protein was preserved in simulations with benzene, and the backbone RMSD of the ECD was similar to control simulations, irrespective of glycoform, indicating that the trimeric assembly of the ECD was unaffected by the presence of benzene (Figure S4). It should be noted that the trimeric coiled coils forming the HR2 domain partially disintegrated as benzene accumulated at its interface (Figure S5), likely because its trimeric interface is primarily composed of an array of hydrophobic residues. Due to this large structural deviation of the HR2 domain, as well as the fact that the HR2 domain is more extensively covered by glycans, we therefore only consider cryptic pockets mapped onto the surface of the ECD of the S protein.

The differences between water-only and benzene simulations revealed multiple cryptic pockets on the surface of the S protein – two of which were previously known from structural studies and have hence been used here as positive controls – and one novel pocket located near the functionally interesting loop encompassing residues 617-628 (Figure 1). We also detected pocket densities near a proposed binding site for bacterial lipopolysaccharide,^27^ but because it is predominantly a surface groove, we have omitted it from further analysis (Figure S6).

**Figure 1.**
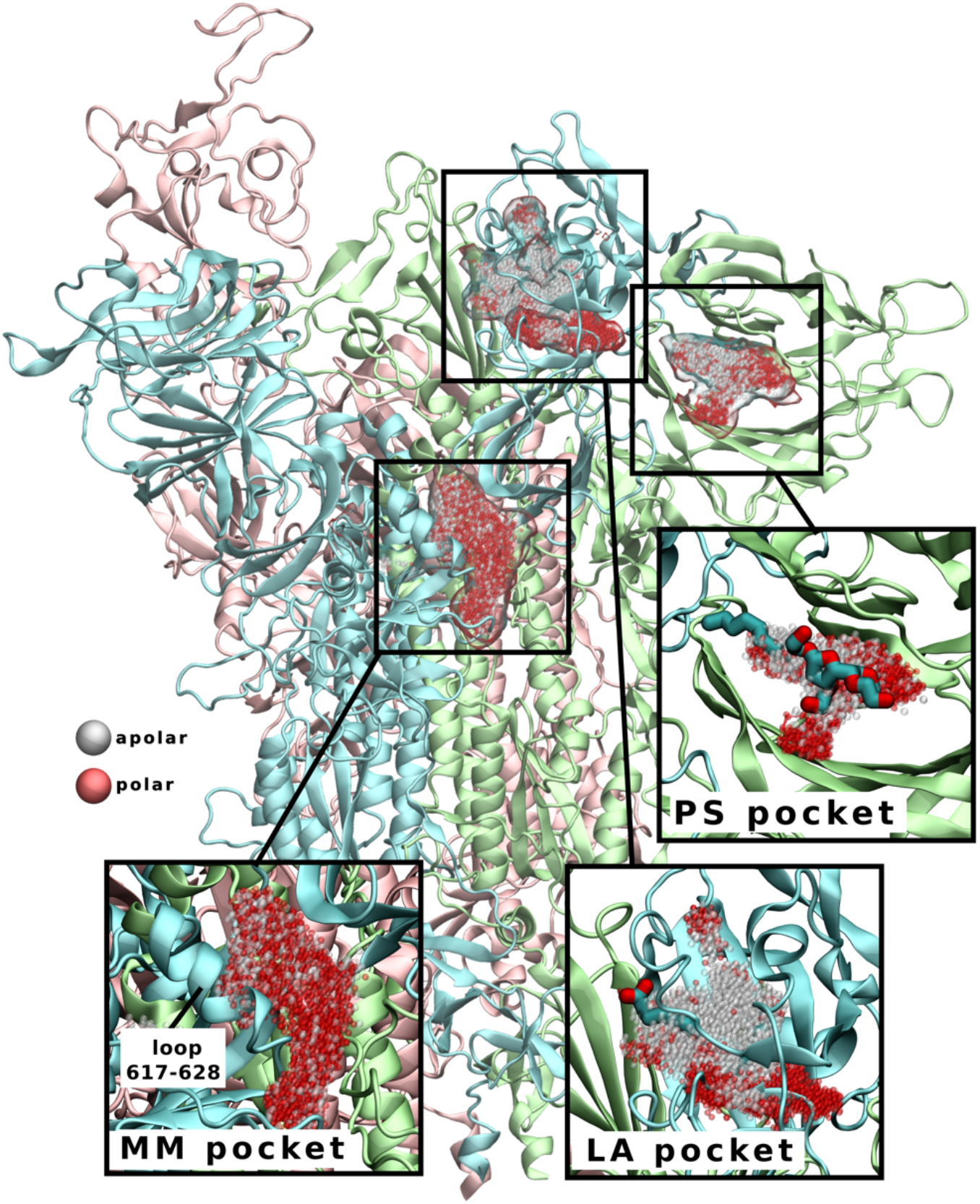
S protein cryptic pockets revealed during simulations in the presence of benzene. Linoleic acid (LA) and polysorbate (PS) pockets are named according to their bound ligands (shown in licorice representation in insets), as their binding sites have previously been identified by cryo-EM.^14–16^ Multimerization (MM) pocket is a surface groove which extends down the interchain interface and underneath the 617-628 loop (labelled in inset). Pockets are shown as clusters of grey (apolar) and red (polar) spheres. The three chains of S protein are coloured as in Figure S1.

A polysorbate 80 (PS) detergent molecule has been observed to bind to the NTD when detergent was present in the formulation of the immunogen.^16^ The hydrophobic tail was embedded in the hydrophobic groove pointing towards the neighbouring chain, with the hydrophilic head more accessible to the protein surface. This site has also recently been shown to bind a haem metabolite, which inhibits access to an antibody epitope on the NTD.^28^ The addition of benzene to the simulation system successfully uncovered the PS-binding pocket, even in the absence of PS. The outline of the mapped pocket was also in agreement with the shape of the hydrophobic portion of the ligand, as confirmed via structural alignment. In comparison, water-only simulations failed to generate pocket densities that matched the shape and properties of the ligand in question, thus demonstrating the power of using hydrophobic probes to reliably induce conformational changes to uncover such cryptic pockets (Figure 2).

**Figure 2:**
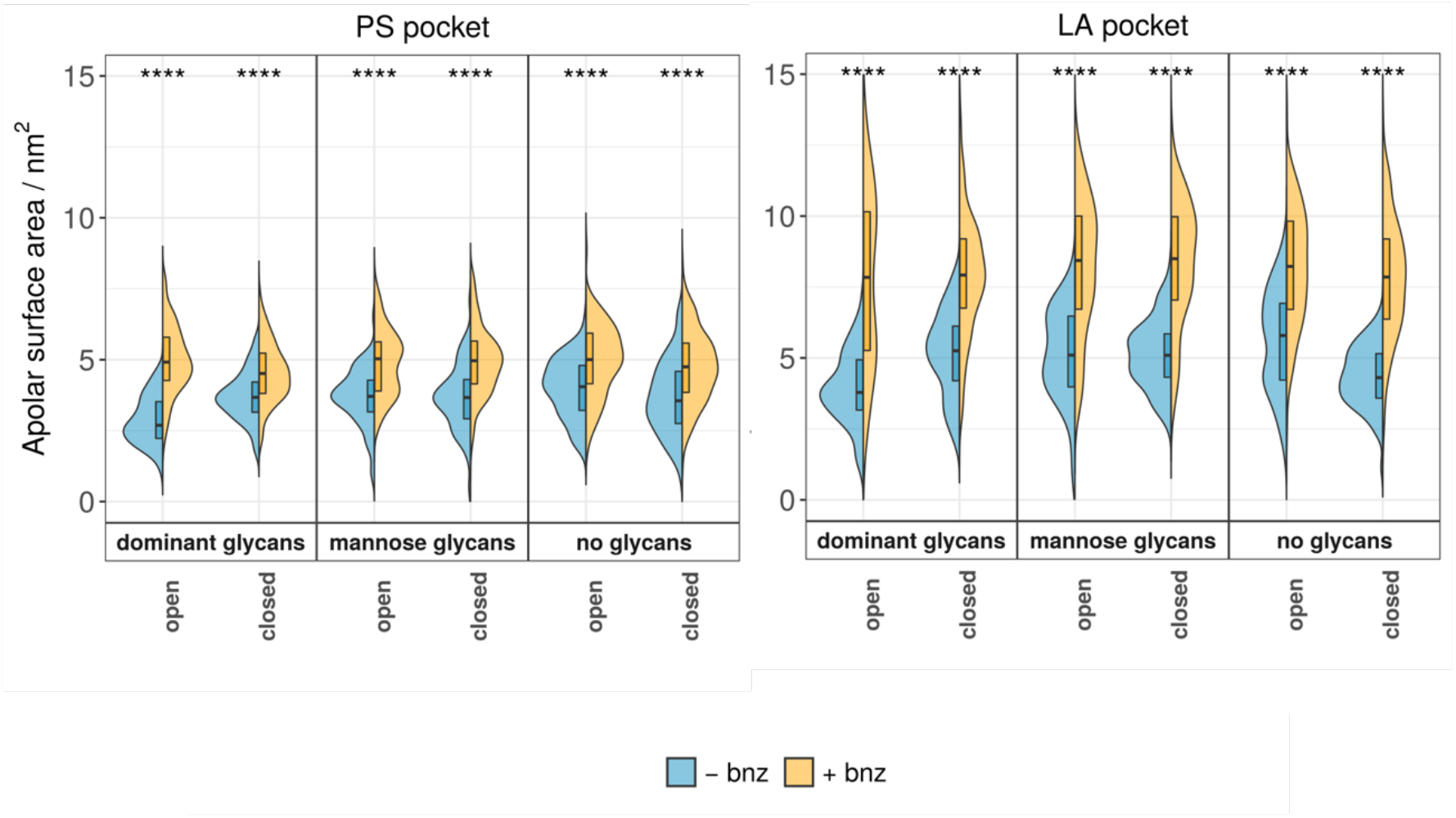
Apolar surface area for positive control polysorbate (PS) and linoleic acid (LA) pockets. “Open” and “closed” systems correspond to S protein chains in the “up” or “down” states, respectively. All three pockets show an increase in their apolar surface area if benzene is present in the system, which indicates that benzene is effective at revealing a larger apolar area of the pocket. *p*-value significance values shown above each plot were calculated using ANOVA (****: p ≤ 0.0001; ***: p ≤ 0.001; **: p ≤ 0.01; *: p ≤ 0.05; ns > 0.05). “Dominant glycans”, “mannose glycans”, and “no glycans” refers respectively to systems in which the S protein has been modelled with either the most dominant species of glycans based on mass spectrometry data^9^, oligomannose-type glycans, or without glycans present.

A second positive control verifying our method was the linoleic acid (LA) binding site, which has been shown to exist in the RBD in multiple cryo-EM structures,^14–16^ but was not present in the structural templates of our initial S models.^3^ We detected increased pocket density in systems with benzene and across all glycoforms. The acyl tail of the LA molecule was accommodated in its entirety in the pocket density, consistent with the nature of the hydrophobic benzene probe, whereas the portion of the pocket outlining the polar carboxylate group was not detected in our simulations. It has been proposed that the presence of LA in the RBD shifts the dynamics of the S protein towards the closed state, whereby all RBDs are in the “down” configuration,^14^ allowing for the interactions of the fatty acid headgroup with the neighbouring chain. As our system was modelled using the S protein ECD in the open state, the arrangement of the RBDs may thus be less able to reproduce the complete outline of the LA binding site that encompasses the fatty acid carboxylate.

Finally, we also detected a novel pocket with a partial cryptic character located on the side of the S protein, which we term the multimerization (MM) pocket. The majority of the pocket volume occupies a shallow surface groove in the interchain region of the S protein, while the cryptic component of the pocket is present in the smaller subsection located underneath the 617-628 loop (Figure 3). Although this short loop is missing from our cryo-EM structural template,^3^ it was predicted to adopt a predominantly helical structure. Additionally, it was shown to be helical in a thermostable disulphide-stabilized S protein construct.^29^ A recent cryo-EM structure of the S protein dimer-of-trimer^16^ shows that this loop is involved in the formation of S protein multimers on the viral surface via its insertion into the NTD of the neighbouring S protein. The cryo-EM map of the S dimer-of-trimer complex shows one 617-628 loop from each trimer interacting with the neighbouring NTD, thus establishing two symmetrical points of contacts between the spikes. Interestingly, when involved in multimerization, the loop is extended and exists as a random coil, instead of the predicted helix.

**Figure 3.**
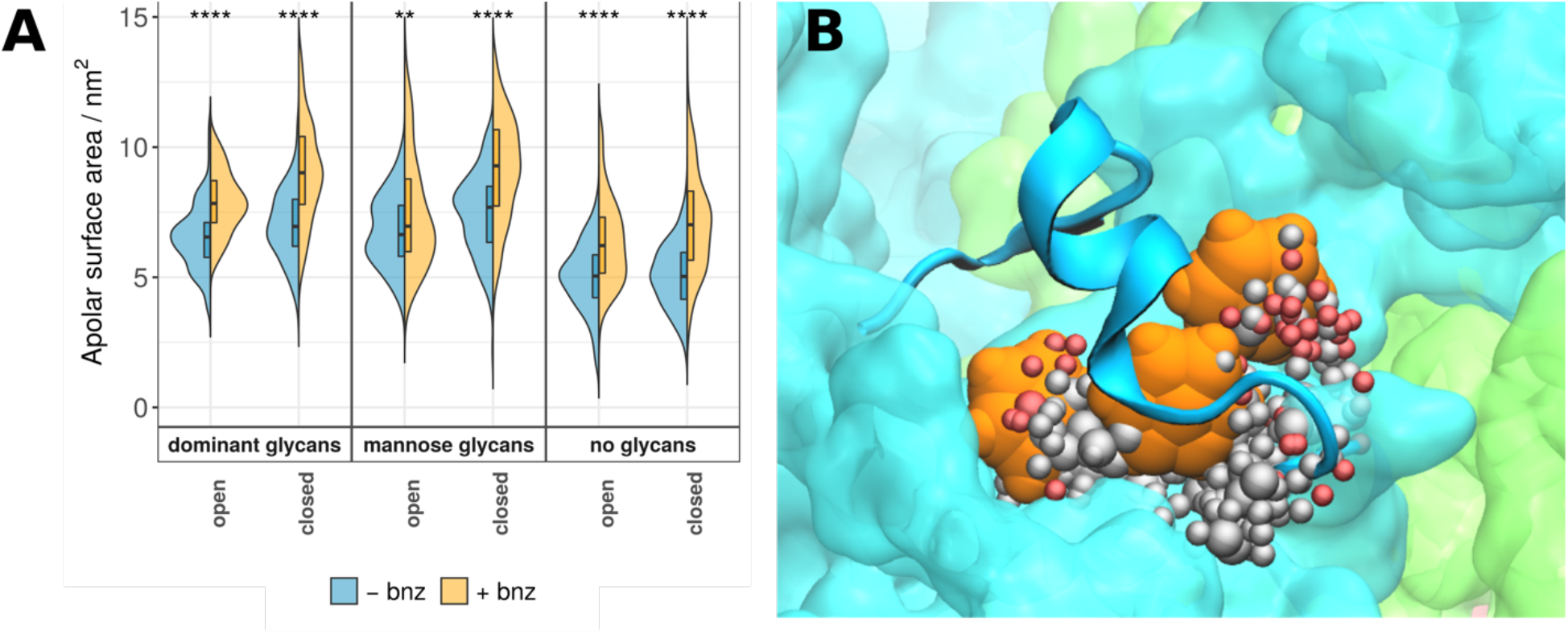
Chemical properties of the MM pocket. **(A)** Apolar surface areas of the MM pocket are shown as violin plots and labelled as in Figure 2. Similar to positive controls, the addition of benzene to the system increases overall apolar component of the pocket in question. System names are as described in Figure 2. **(B)** A portion of the MM pocket packed underneath the 617-628 loop. Three benzene molecules (in orange) are able to interact with the protein surface and occupy a predominantly hydrophobic binding pocket (spheres representing the pocket are colour coded as in Figure 1).

We then examined the behaviour of the loop via HDX-MS experiments (see Methods). Peptides encompassing the loop exhibited a multimodal spectrum that reflects conformational ensemble behaviour or intratrimer conformational heterogeneity, with a major lower exchanging population and minor, higher exchanging population in solution (Figure 4, Figure S7).^30,31^ We compared this with our simulated systems and observed a similar multimodality in the behaviour of the loop, or in this case, its associated peptide. Solvent accessibility of the simulated peptide reflected a similar pattern of behaviour, with a large population of states with lower solvent accessibility, and rarer occurrences of states with a more accessible peptide surface. Cluster analysis similarly shows that the peptide is in a helical state for ∼90% of cumulative simulation time, but occasionally rearranges itself into less structured random coil states (Figure S8A).

**Figure 4.**
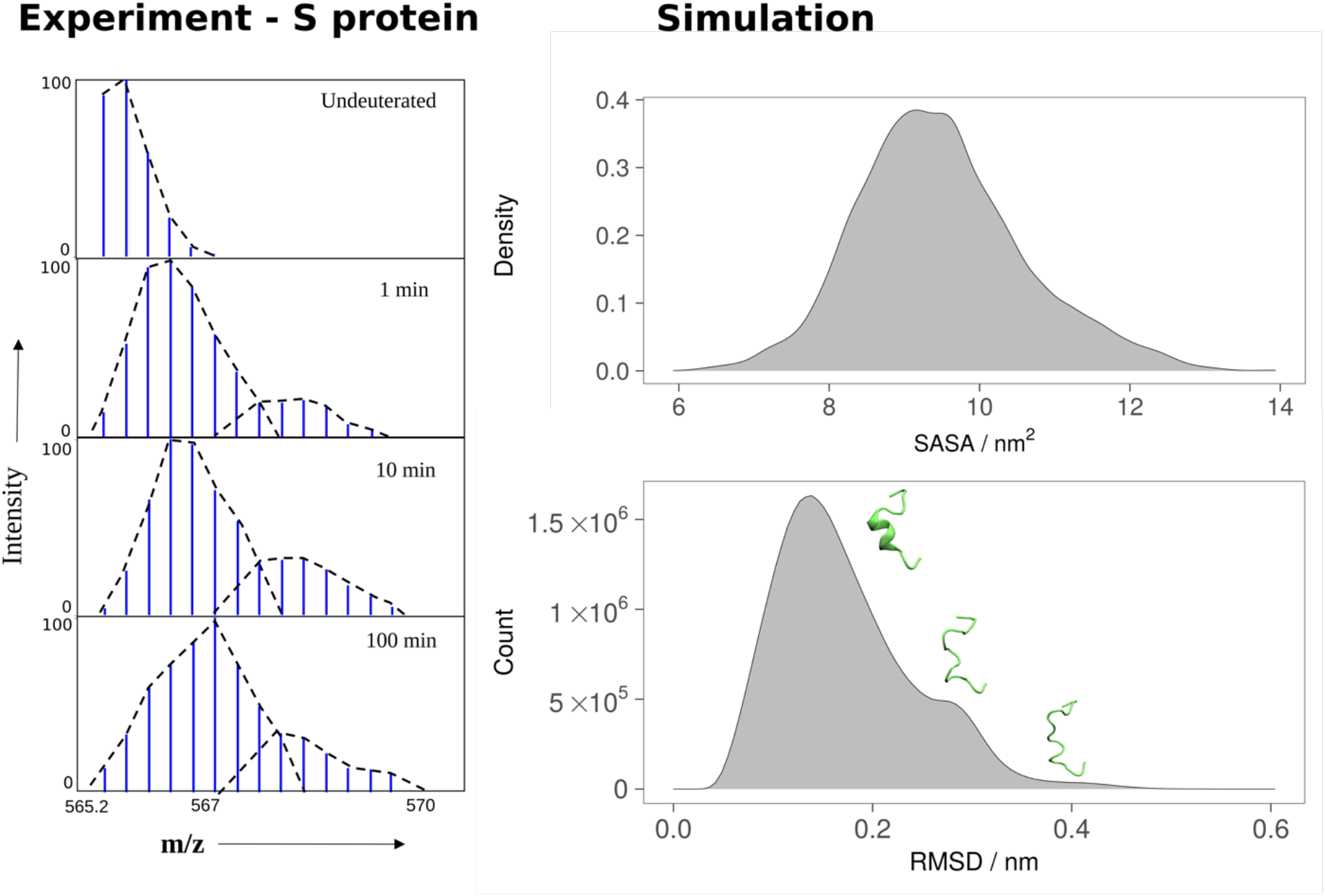
The dynamic properties of peptide spanning residues 617-632. HDX-MS mass spectrum of the peptide at labelled time points (left) shows a bimodal profile with a minor higher exchange population, indicating a conformational ensemble property in solution. In MD simulations, the loop containing this peptide exhibits a similar bimodal characteristic as assessed via solvent-accessible surface area (SASA) and conformational cluster analysis (right).

The observed conformational heterogeneity in loop behaviour, both in experiment and in simulation, may be of functional importance for the S protein, particularly in the context of higher order S protein complex formation. Consistently, mutations in the loop have been shown to reduce infectivity and expression of SARS-CoV-2, indicating a potential role for the loop in viral assembly.^16^ In our simulations, the MM pocket behind the 617-628 loop interacted with one, two, or rarely three benzene molecules at any given time (Figure 3B and S8B). A multiple sequence alignment indicates that the surface of the MM pocket is well-conserved across different coronaviruses (Figure S9). Interestingly, the pocket also contains residue D614 which, when mutated to glycine, results in faster viral transmission, more efficient infection and replication,^32^ and higher S protein density on the viral surface.^33^ Additionally, the pocket is within close proximity to residues A570 and H655, which are mutated in the novel SARS-CoV-2 strains B.1.1.7 and B.1.1.28, respectively, both of which are associated with increased transmissibility and severity of infection (Figure S10).^34^ Thus, the predominantly hydrophobic nature of the pocket, its potential to interact with aromatic moieties commonly present in the drug molecules, its well conserved surfaces and, most importantly, its proximity to the functionally relevant 617-628 loop and mutated residues in novel SARS-CoV-2 variants with higher transmission rate, are all promising indicators of the potential druggability of the MM pocket.

In summary, we have demonstrated the power of the benzene mapping technique to delineate cryptic hydrophobic pockets that are of interest for drug and monoclonal antibody development targeting the SARS-CoV-2 S glycoprotein. In addition to successfully reproducing two independently identified cryptic pockets, we also uncovered a novel and potentially druggable pocket on the S protein ECD surface. Simulations of different S glycoforms indicated that cryptic pockets are present in all glycosylation states, suggesting that these pockets could be targeted in all stages of viral maturation. Complementing efforts in antibody-based therapy and vaccine development, rational design of small-molecule drugs targeting S protein pockets may provide an essential therapeutic practice for combatting the COVID-19 pandemic.

## ASSOCIATED CONTENT

### Supporting Information

Methodological details, list of simulations, system validation, additional simulation data.

### Funding sources

No competing financial interest has been declared.

## ACKNOWLEDGEMENTS

This work was supported by BII of A*STAR. Simulations were performed on the petascale computer cluster ASPIRE-1 at the National Supercomputing Centre of Singapore (NSCC), the A*STAR Computational Resource Centre (A*CRC), Iridis 5 supercomputer at the University of Southampton and ARCHER provided by the UK HECBioSim.

---

**For Table of Contents Only**

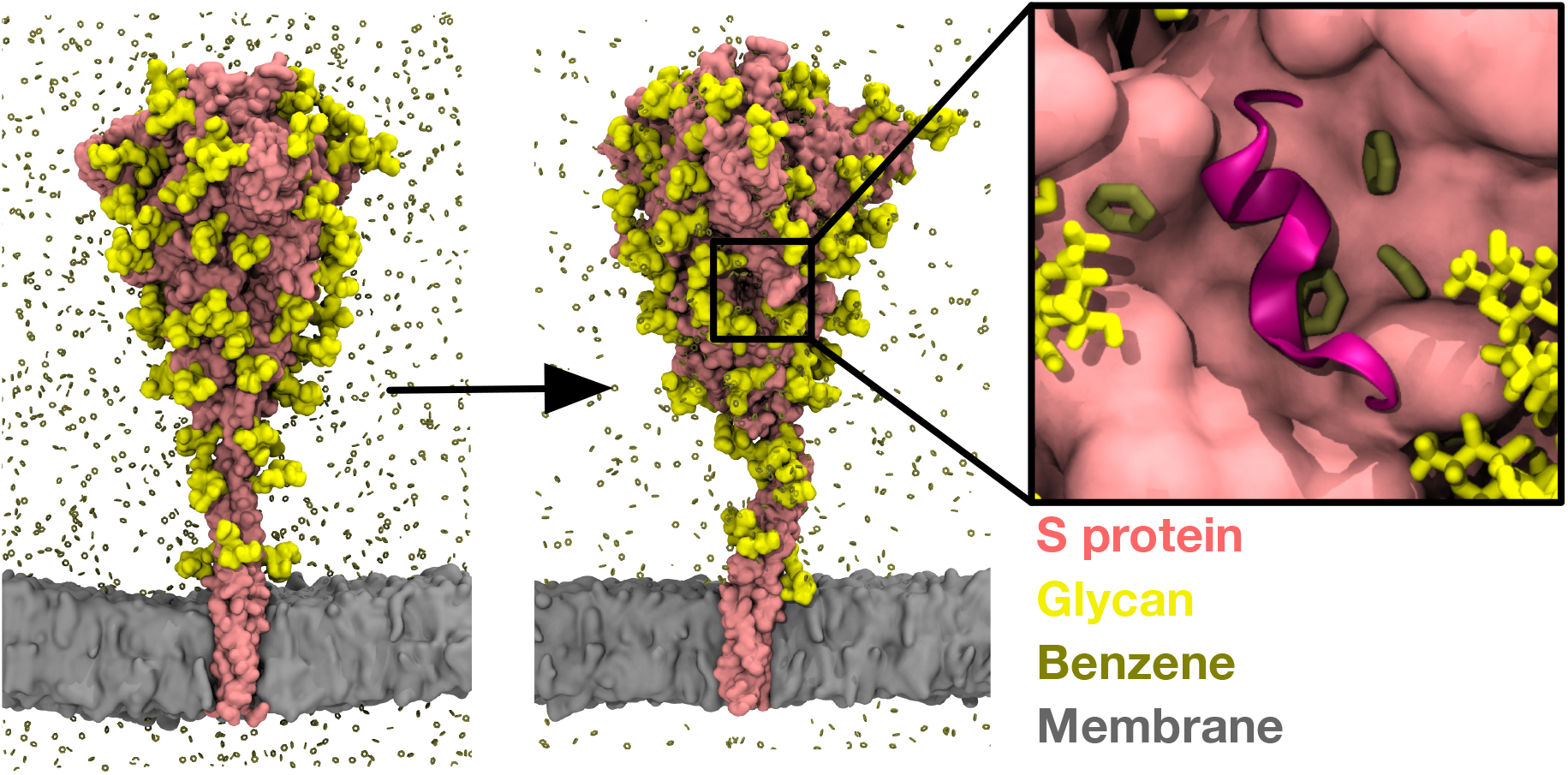

