## Supplementary Material for "Uncovering cryptic pockets in the SARS-CoV-2 spike glycoprotein"

[†]Joint first authors

*To whom correspondence should be addressed:

### METHODOLOGICAL DETAILS

#### Integrative Modelling of the SARS-CoV-2 S Glycoprotein

Modeller version 9.21[1] was used to construct homology models of full-length S glycoprotein. The full sequence was obtained from the complete genome of SARS-CoV-2 (GenBank: MN908947).[2] The cryo-EM structure of the S protein ECD in the open state (PDB: 6VSB)[3] was used as the main template for the ECD.[3,4] Missing loops in the RBD up state were modelled using the cryo-EM structure of SARS-CoV-2 RBD bound to the angiotensin converting enzyme (ACE) 2 receptor (PDB: 6M17),[5] while missing loops in the N-terminal domain (NTD) and the C-terminal of the ECD were modelled using the cryo-EM structure of S ECD in the closed state resolved at a higher resolution (PDB: 6XR8).[6] The heptad repeat 2 (HR2) domain was modelled based on the NMR structure of SARS-CoV HR2 domain (96% sequence identity) in the prefusion conformation (PDB: 2FXP).[7] To date, there is no structural information for the transmembrane (TM) domain of the S protein from any coronaviruses. To estimate the position of the TM domain, the PSIPRED secondary structure prediction web server was used.[8] The server predicted residue 1213-1237 to be within the TM helix. The presence of a GxxxG motif within this sequence suggests that the three helices of the S protein subunits form an oligomeric assembly in the membrane.[9] We therefore used the putative TM domain sequence to search for a homotrimeric TM structure with reasonable sequence similarity. The NMR structure of human immunodeficiency virus 1 (HIV-1) gp-41 TM domain (PDB: 5JYN),[10] which shares 27% sequence identity, was subsequently used as template. Ten models were built and the three models with the lowest discreet optimized protein energy (DOPE) scores[11] were chosen for further stereochemical assessment using Ramachandran analysis.[12] The model with the lowest number of outlier residues was subsequently selected for further modifications. The model with the lowest number of outlier residues was subsequently selected for further modifications.

Two post-translational modifications were incorporated into the models: palmitoylation and glycosylation. Palmitoylation of two cysteine clusters in the SARS-CoV S protein has been shown to be important in membrane fusion with the host cell.[13] We therefore added palmitoylation to cysteine residues at position 1236, 1240 and 1243 found in these two membrane proximal clusters. There are 22 N-glycosylation sites on each subunit of the S proteins, which likely play a crucial role in immune evasion by blocking access to protein epitopes. For each model, we built three glycoforms: i) with the most dominant glycan found on each site based on mass spectrometric data[14]; ii) with an oligomannose-type glycan ($Man_9GlcNAc_2$) on all sites to represent the unprocessed glycosylated protein; and iii) without glycans. Palmitoylation and glycosylation of the S protein models was performed using the CHARMM-GUI Glycan Reader and Modeller web server.[15]

**Building Model Membrane**

A 25 x 25 $nm^2$ patch representing the endoplasmic reticulum-Golgi intermediate compartment (ERGIC) membrane was built using the CHARMM-GUI Membrane Builder web server.[16] The membrane is symmetric and the composition is 47% phosphatidylcholine (PC), 20% phosphatidylethanolamine (PE), 11% phosphatidylinositol-phosphate (PIP), 7% phosphatidylserine (PS) and 15% cholesterol.[17–19] All of the lipid tails were modelled as 1-palmitoyl-2-oleoyl (PO). Energy minimization and equilibration were performed according to the CHARMM-GUI protocols.

**Setup and Molecular Dynamics Simulations of Benzene-free Systems**

The CHARMM36 force field was used to parametrize the full-length S glycoprotein models.[20] The protein was inserted into the model membrane based on the position of the TM domain. Overlapping lipid residues were removed and the steepest descent method was performed to minimize the system. The system was solvated with TIP3P water molecules and neutralized with 0.15 M NaCl salt. Step-wise minimization and equilibration adapted from CHARMM-GUI standard protocols were performed.[21] During equilibration, the temperature of the system was maintained at 310 K using the Berendsen thermostat with a time constant of 1 ps, while the pressure was kept at 1 atm by a semi-isotropic pressure coupling using the Berendsen barostat with a time constant of 5 ps.[22] The smooth particle mesh Ewald (PME) method[23] with a real-space cut-off of 1.2 nm was utilised to calculate the electrostatic interactions, whereas the Van der Waals interactions were truncated at 1.2 nm and the force switch smoothing function applied between 1.0 to 1.2 nm. An integration time step of 1 fs and 2 fs were used at the early and late steps of equilibration, respectively, with the LINCS algorithm utilised to constrain all covalent bonds with hydrogen atoms.[24] After equilibration, 200 ns production simulations were conducted. The Nosé-Hoover thermostat with a time constant of 1 ps was used to maintain the temperature,[25,26] while the Parrinello-Rahman barostat with a time constant of 5 ps was used to maintain the pressure.[27] A 2-fs integration time step was employed during this run.

**Setup and Molecular Dynamics Simulations of Benzene Systems**

A protocol for setting up benzene probe simulations in membrane systems was adapted from our previous work.[28] Benzene molecule partial charges were adjusted from phenylalanine aromatic ring parameters defined in CHARMM36 force field[20] so that the distribution of charges was uniform across all six carbon atoms. A virtual site was added at a geometric centre of each benzene molecule and it served as a point of repulsion between the benzene probe and the membrane. As the membrane composition differed from the one used in the original paper, repulsion point positions and the Lennard-Jones σ-value had to be adjusted to be effective for the ERGIC membrane composition. A repulsion point was placed on a carboxyl oxygen (O22/O32), one in each fatty acid tail. Repulsion point atoms were present in all lipid types (including protein-bound palmitoyl group), except for cholesterol. To account for increased gaps between membrane repulsion points due to the presence of cholesterol, the σ-value for the interactions between benzene dummy atoms and membrane repulsion points was increased to 1.4 nm. The setup was tested for the effects of aggregation and probe sequestration on a small ERGIC membrane (7 nm × 7 nm) and during a 100 ns simulation, benzene molecules remained solvated and outside the bilayer.

Benzene probes were added to a simulation box in a 0.2 M concentration. A lower probe concentration allowed for the omission of benzene-benzene repulsions and exclusions, as the probes were less likely to aggregate.[29] Subsequently, the rest of the system was solvated with TIP3P water and 0.15 M NaCl salt. Minimization, equilibration, and production simulations followed the same protocol as for the benzene-free systems. All simulations were performed with GROMACS 2018[30] and the list of simulations is provided in Table 1.

**Hydrogen-Deuterium Exchange Mass Spectrometry of the S Protein**

Deuterium exchange mass spectra of the peptides 617-632 and 621-633 are reported from the data deposited in ProteomeXchange Consortium via the PRIDE partner repository[31] with dataset identifier: PXD23138 and reported by Palur et al.[32] Briefly, Deuterium exchange reaction was performed by incubating purified recombinant S protein (PBS, pH 7.4) in PBS buffer containing 90% $D_2O$ at 37°C for 1, 10 and 100 minute of labelling time. The deuterium exchange reaction was stopped by mixing the reaction mixture with prechilled quenched buffer (1.5 M GnHCl and 0.25 M Tris(2-carboxyethyl) phosphine-hydrochloride (TCEP-HCl)) to lower the pH to 2.4 and incubated in ice for 1 minute before online pepsin digestion and mass spectrometry analysis. Quenched samples were injected into nanoUPLC HDX sample manager (Waters, Milford, MA). Immobilized Waters Emzymate BEH pepsin column (2.1 × 30 mm) was used to perform online digestion in 0.1% formic acid in water at 100 μl $min^{-1}$ flow rate. Pepsin-proteolyzed peptides were trapped in a 2.1 × 5 mm C18 trap (ACQUITY BEH C18 VanGuard Pre-column, 1.7 μm, Waters, Milford, MA). Elution of trapped peptides was performed using acetonitrile gradient of 8% to 40% in 0.1% formic acid at flow rate 40 μl $min^{-1}$ into reverse phase column (ACQUITY UPLC BEH C18 Column, 1.0 × 100 mm, 1.7 μm, Waters) pumped by nanoACQUITY Binary Solvent Manager (Waters, Milford, MA). Electrospray ionisation mode was used to spray ionised peptides and HDMSE mode of detection was implemented on SYNAPT G2-Si mass spectrometer (Waters, Milford, MA). 200 fmol $μl^{-1}$ of [Glu1]-fibrinopeptide B ([Glu1]-Fib) is injected for lock spray correction at a flow rate of 5 μl $min^{-1}$. Protein Lynx Global Server (PLGS v3.0, HDMSE mode) was used to identify the mass spectra of undeuterated protein samples on a separate sequence database of each protein sequence. Further, peptides were filtered using minimum intensity cutoffs of 2500 for product and precursor ions, precursor ion mass tolerance of <10 ppm and minimum products per amino acids of 0.2 using DynamX v 3.0 (Waters, Milford, MA). All the experiments were performed in triplicate and not corrected for back exchange.

**Analysis**

Protein, sugar and membrane structural properties were analysed using GROMACS tools, MDAnalysis[33] and VMD.[34] Pockets on the surface of the S protein ectodomain were analysed using MDPocket.[35] Initial pocket mapping of the entire ectodomain structure was followed by repeated mapping and property characterization of each individual pocket of interest. Pockets were analysed across all simulated trajectories in 5 ns snapshot intervals. Sequence conservation of the S protein was analysed using the ConSurf webserver.[36] The HMMER algorithm[37] was used to pull 40 unique sequences with 35-95% sequence identity to the SARS-CoV-2 S protein (GenBank: MN908947)[2] from the UniProt database.[38] Multiple sequence alignment was built using the MAFFT program.[39]

**Table S1: List of simulations**

| S protein glycosylation state | Benzene concentration (M) | Simulation time (ns) | Number of replicates |
|---|---|---|---|
| Non-glycosylated | 0 | 200 | 2 |
| High mannose | 0 | 200 | 2 |
| Dominant glycans | 0 | 200 | 2 |
| Non-glycosylated | 0.2 | 200 | 3 |
| High mannose | 0.2 | 200 | 3 |
| Dominant glycans | 0.2 | 200 | 3 |

### SUPPLEMENTARY FIGURES

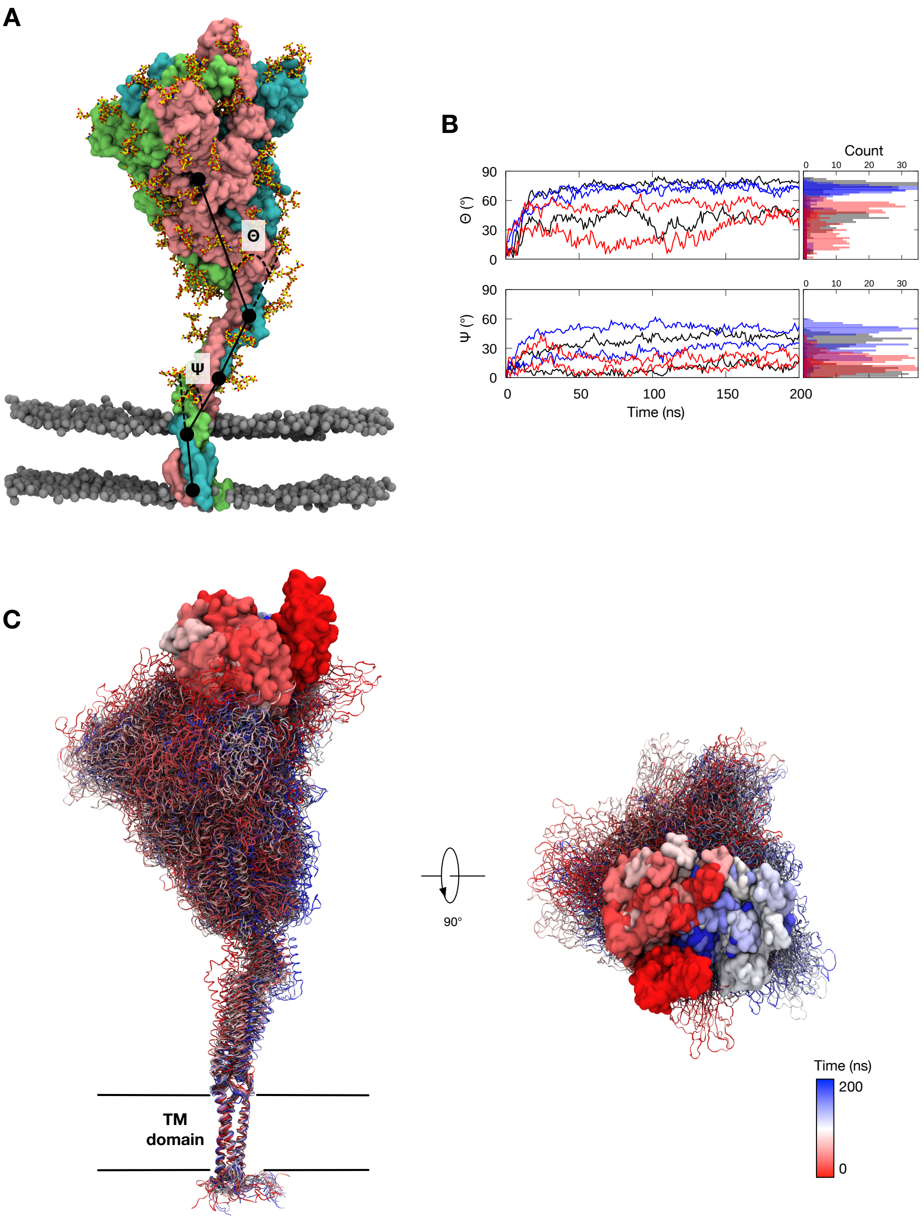


**Figure S1: S protein bending dynamics during simulations without benzene. (A)** A snapshot from the end of a simulation of S protein open conformation with dominant glycans. The three protein subunits are shown in surface representation and coloured cyan, pink and green; the glycan moieties are shown in stick representation and coloured yellow; the lipid phosphorus atoms are shown in sphere representation and coloured grey. **(B)** Top, bending angle, Θ, measured as the acute angle between the centres of mass of the ECD (residue 27-1141), the hinge region (residue 1142-1156) and the HR2 domain (residue 1157-1207). Bottom, bending angle, Ψ, measured as the acute angle between the HR2 domain, the hinge region (residue 1208-1212) and the TM domain (residue 1213-1239). Black, dominant glycans; blue, mannose glycans; red, no glycans. (C) Snapshots of S protein at 10 ns intervals from a simulation, fitted onto the TM domain fixed. The RBD is highlighted in surface representation. Glycan molecules are omitted for clarity.

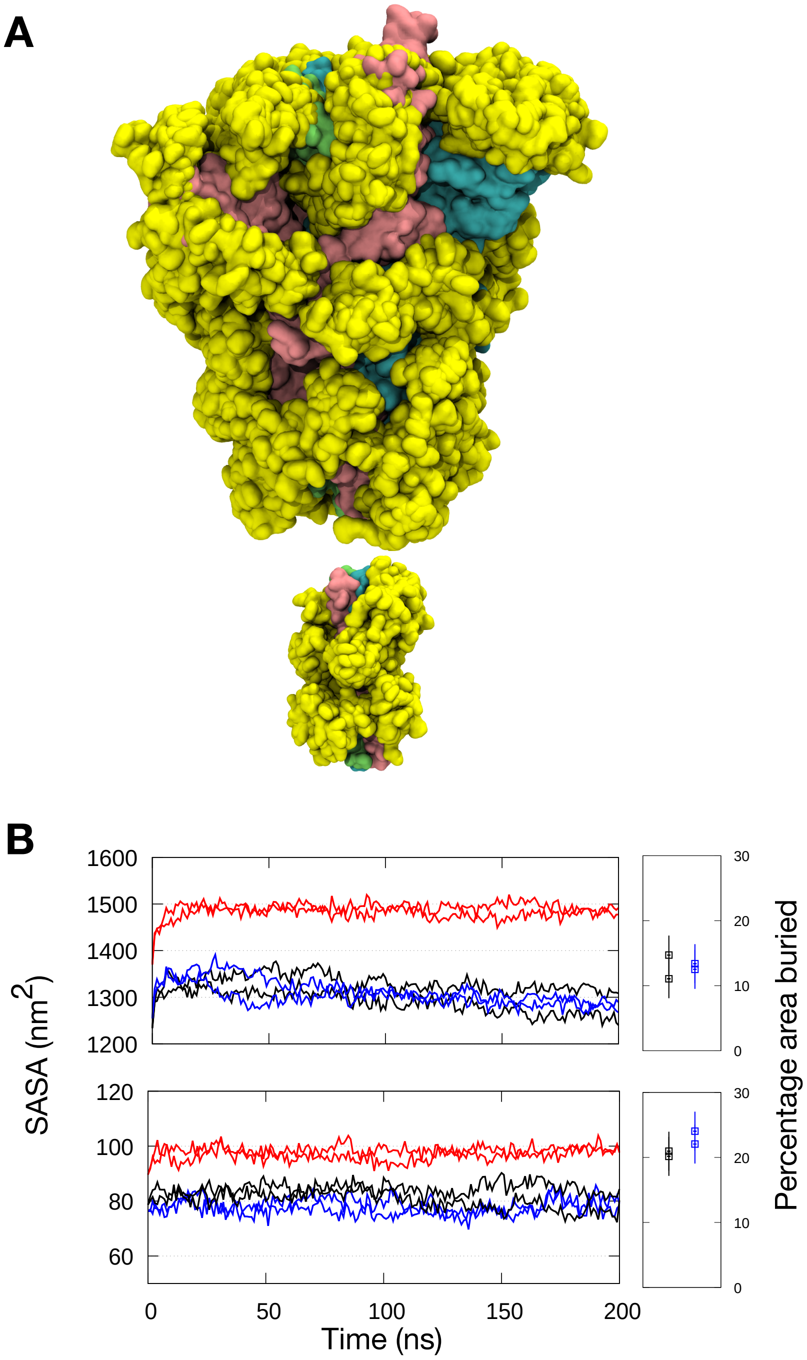


**Figure S2: S protein shielding by glycans during simulations without benzene. (A)** Snapshots of S protein ECD (top) and stalk (bottom) coloured cyan, pink and green with glycans in yellow at 10 ns intervals from a simulation of S protein with dominant glycans. **(B)** Solvent accessible surface area (SASA) of the ECD (top) and the stalk (bottom). Black, dominant glycans; blue, mannose glycans; red, no glycans. Percentage of surface area buried by glycans is shown on the right, average from the last 50 ns of the simulations with the error bars showing standard deviations along the trajectories. The radius of solvent probe used for calculation was 0.14 nm.

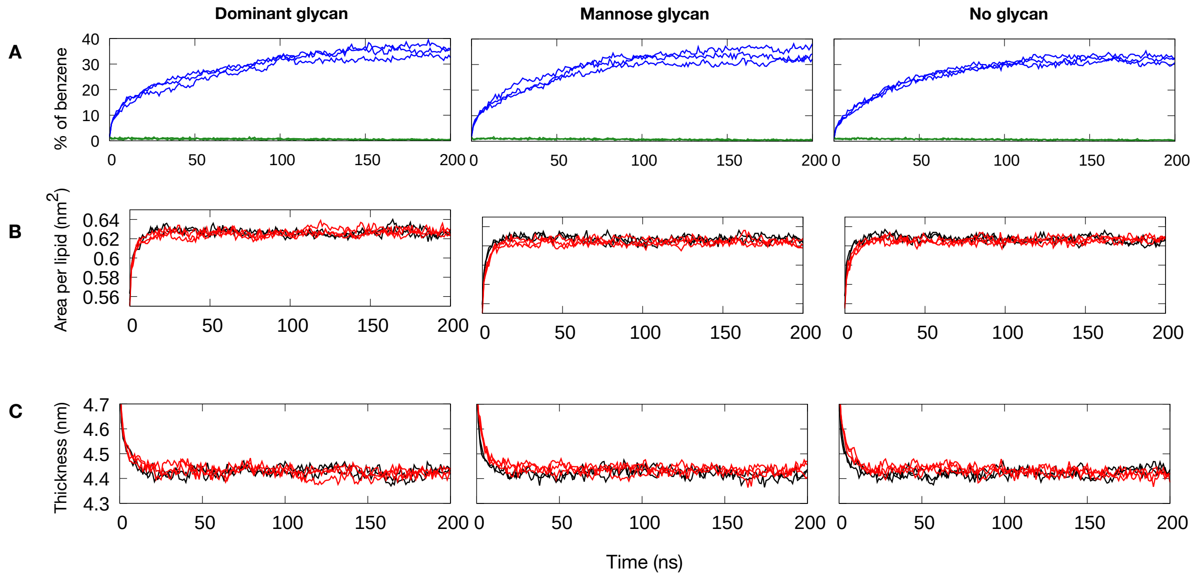


**Figure S3: Membrane structural properties during simulations of S protein with and without benzene. (A)** Percentage of benzene in contact with the membrane lipids (green) or S protein (blue) throughout the simulations with benzene. Cut-off for contact is 0.4 nm. **(B), (C)** Area per lipid and membrane thickness in simulations with (red) and without benzene (black).

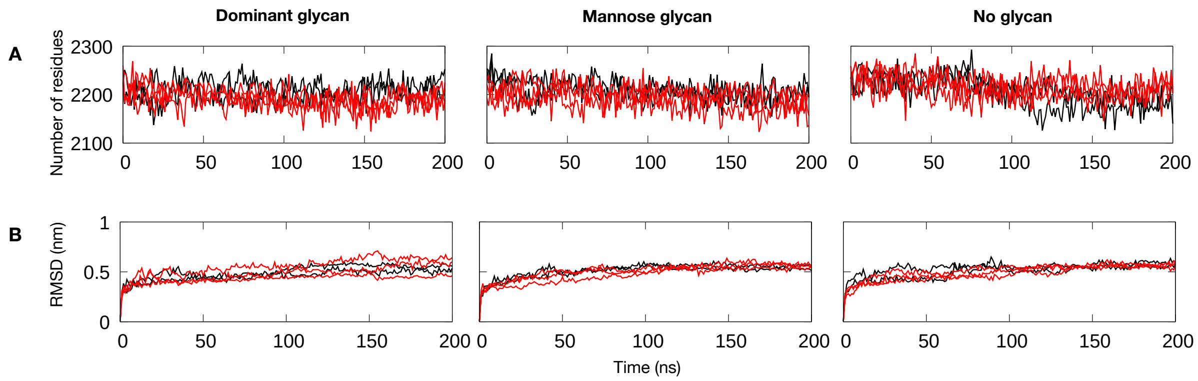


**Figure S4: Protein structural properties during simulations of S protein with and without benzene. (A)** Total number of residues with secondary structural elements (α-helix, β-sheet, β-bridge and turn). **(B)** Backbone RMSDs for the ECD (residue 27-1146). Data correspond to simulations with (red) or without benzene (black).

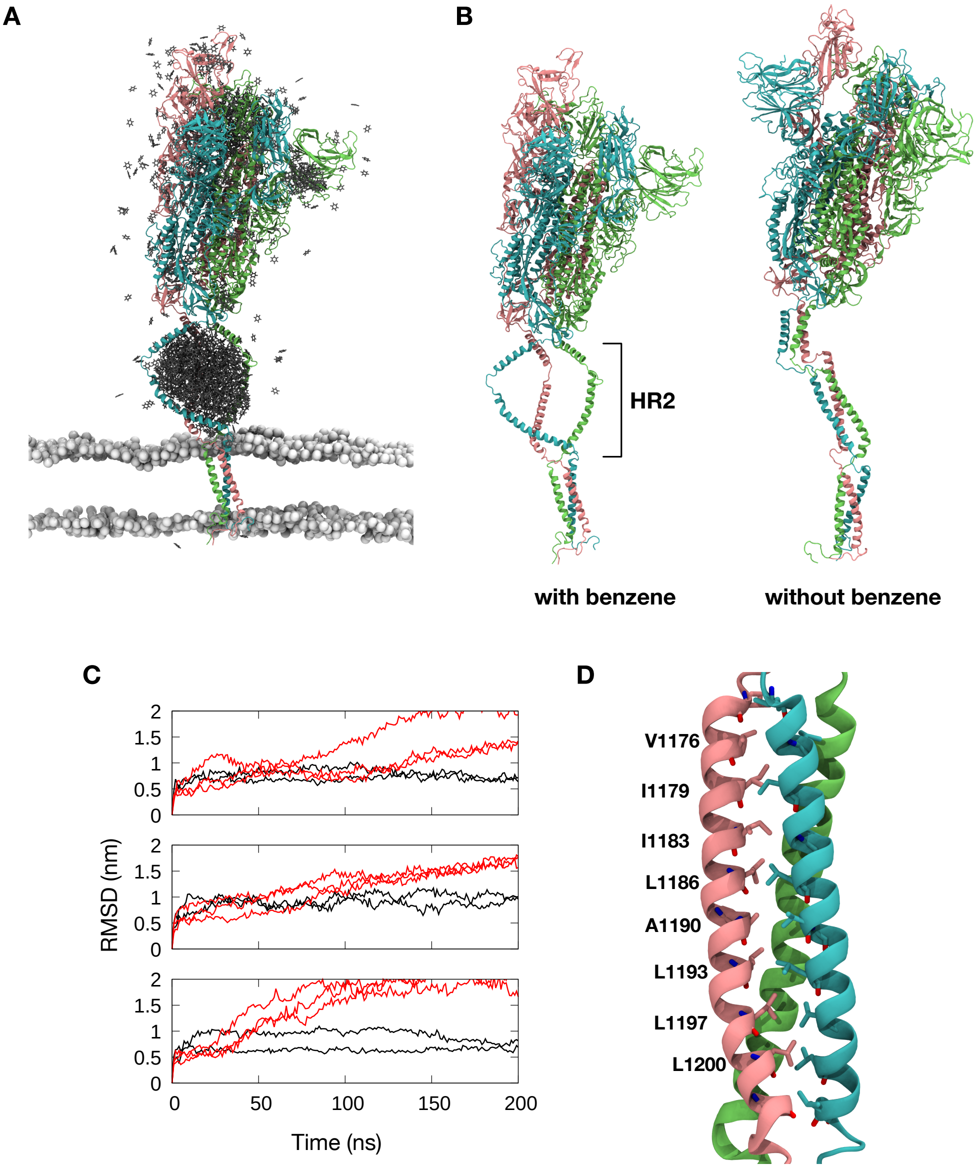


**Figure S5: Aggregation of benzene around the HR2 domain. (A)** The final snapshot from one of the simulations of glycosylated S protein open state with 0.2 M benzene. Benzene is shown in stick representation and coloured grey. For clarity, only benzene found within 2.0 nm of the protein is shown. **(B)** Comparison of protein structures at the end of simulations with and without benzene. HR2 domain is labelled. **(C)** Backbone RMSDs for the HR2 domain from simulations with (red) and without benzene (black). Dominant glycans (top); mannose glycans (middle); no glycans (bottom). **(D)** Hydrophobic residues at the interface of the HR2 domain.

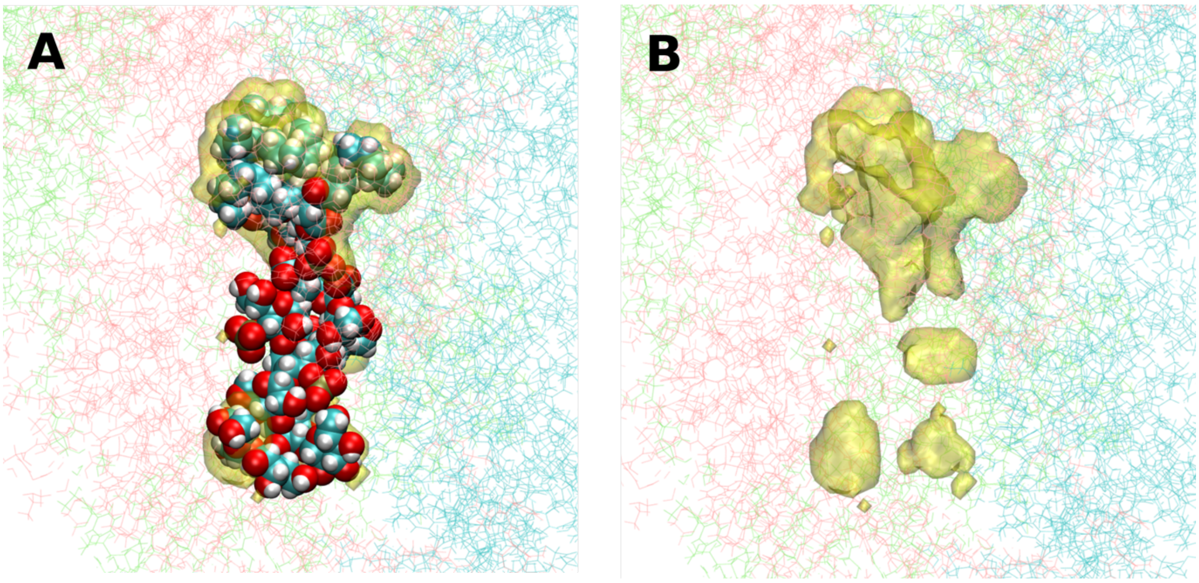


**Figure S6: Location and properties of proposed LPS pocket.** LPS pocket is defined as any pocket area within 0.2 nm of the LPS molecule. **(A)** LPS molecule shown in spheres docked in a proposed LPS pocket, based on Petruk et al.[40] Mapped pocket area is shown as yellow surface. LPS pocket predominantly outlines the fatty acid tail portion of the LPS, while the hydrophilic sugar components are largely outside any detected pocket regions. **(B)** The outline of the LPS pocket without a bound ligand.

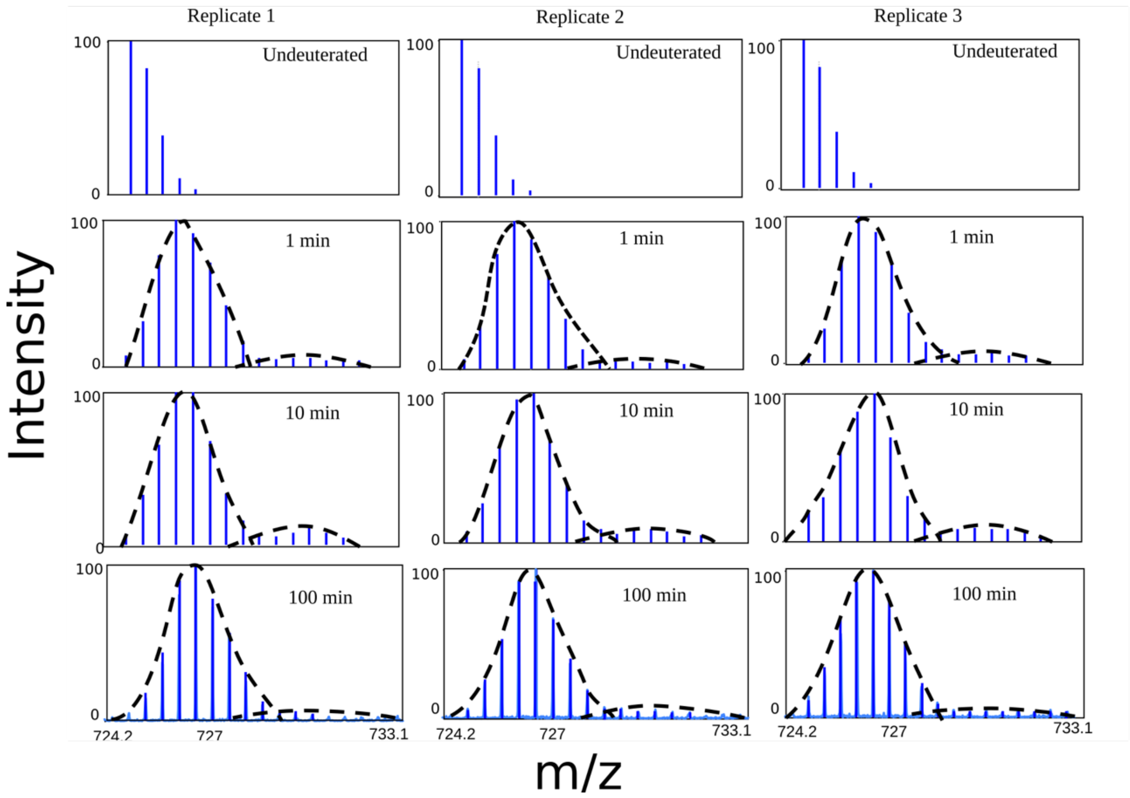


**Figure S7: HDX-MS mass spectra of the overlapping 621-633 peptide.** Mass spectra (blue) of 621-633 peptide from three replicates demonstrates EX1 mode (black dashed lines) of deuterium exchange kinetics at all labelling times, similar to exchange for 617-632 peptide.

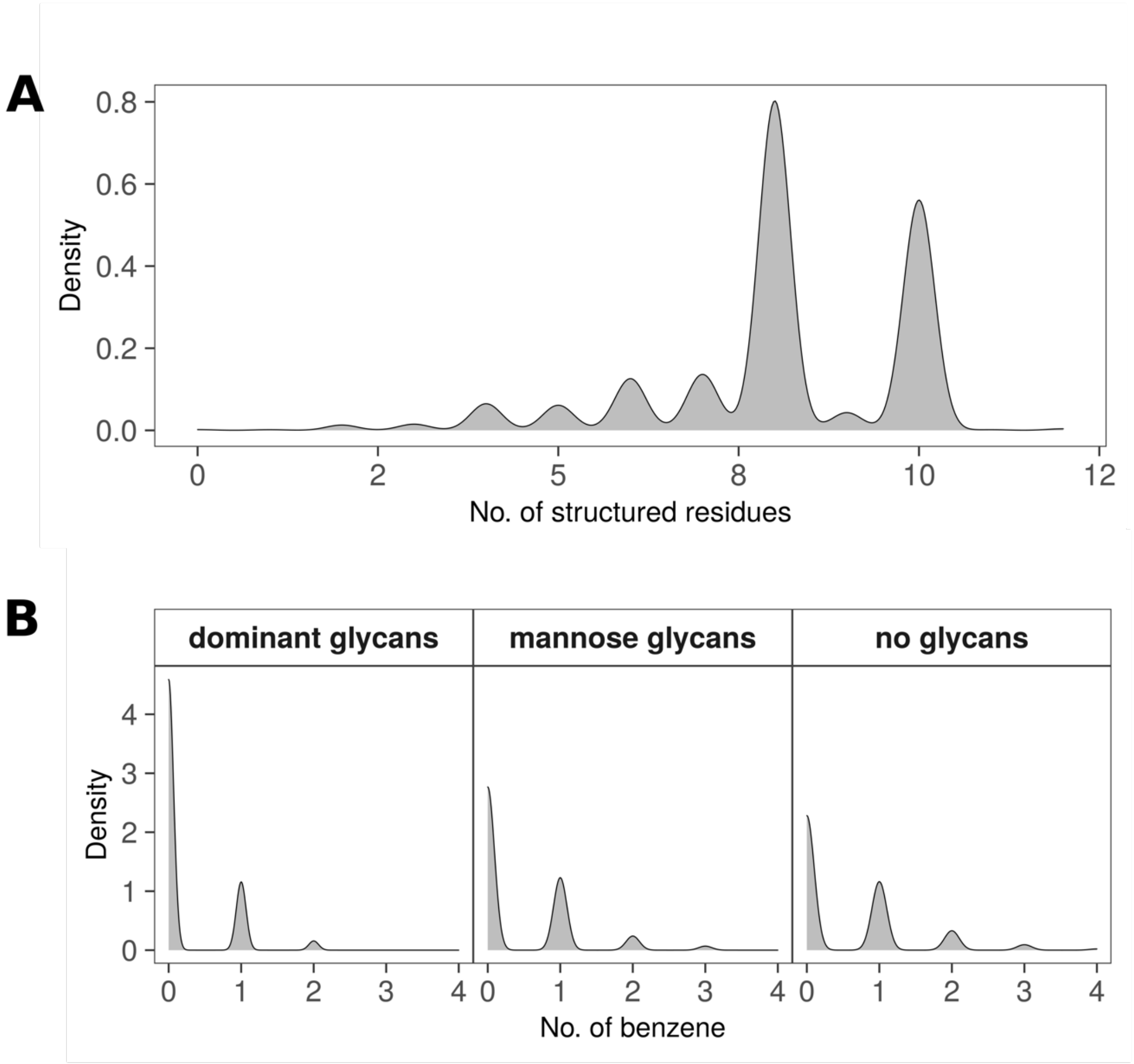


**Figure S8: Properties of MM pocket segment. (A)** Cumulative density distribution of number of residues in the 617-628 loop that are structured (either as $\alpha$-helix or turn). **(B)** Number of benzene molecules that occupy a segment of the MM pocket underneath the 617-628 loop (see Figure 4B). The maximum number of benzenes occupying the pocket at any given point was three.

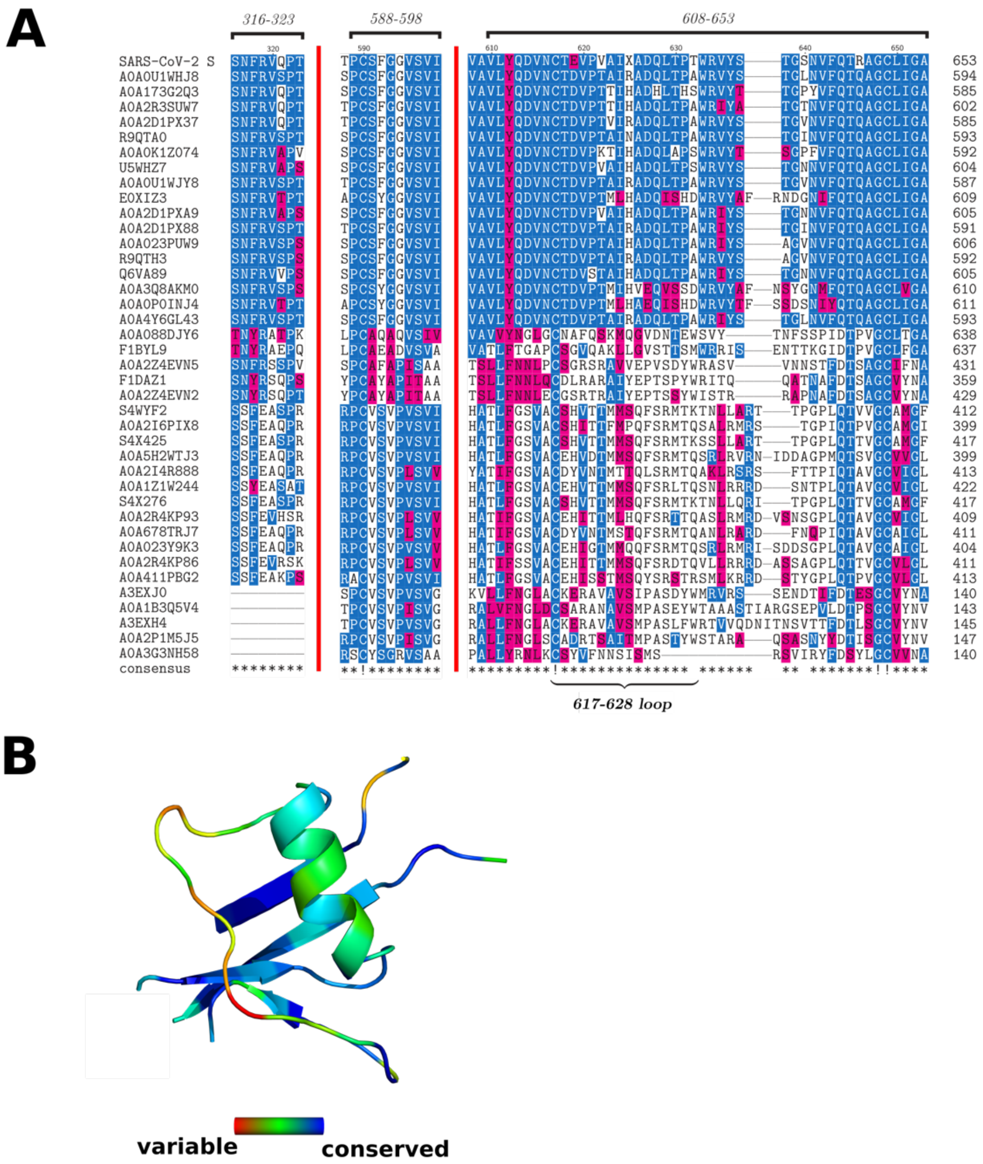


**Figure S9: Conservation of the MM pocket segment underneath the 617-628 loop.** Sequences were selected and scored using CONSURF webserver (details in Methods). **(A)** Multiple sequence alignment of the MM pocket segment. Residues are shaded based on similarity, using a threshold of 30%. **(B)** MM pocket segment is shown in cartoon representation and is coloured according to CONSURF score.

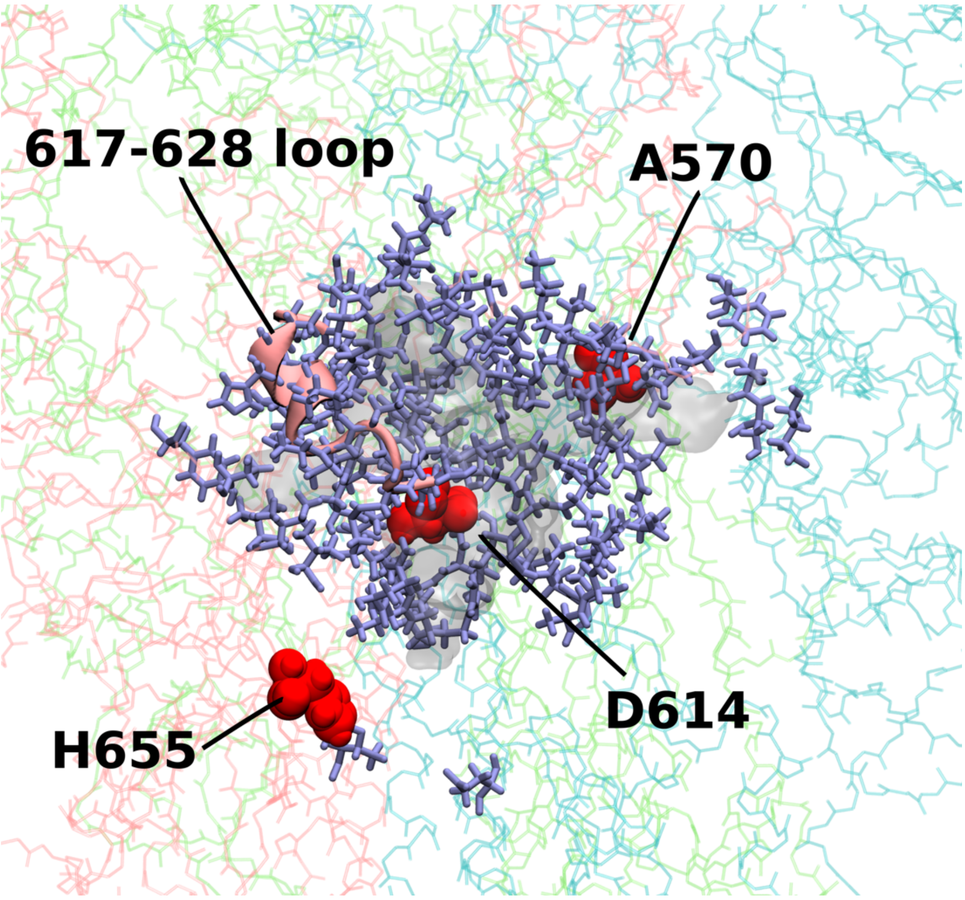


**Figure S10: MM pocket and its associated residues.** The pocket is shown in transparent surface representation, residues surrounding the pocket are shown in licorice, and the 617-628 loop in cartoon. The remainder of the S protein is shown in pink, cyan, and lime lines. Residues close to the MM pocket and mutated in novel SARS-CoV-2 strains are shown in red spheres.
